# United States wildlife and wildlife product imports from 2000-2014

**DOI:** 10.1101/780197

**Authors:** Evan A. Eskew, Allison M. White, Noam Ross, Kristine M. Smith, Katherine F. Smith, Jon Paul Rodríguez, Carlos Zambrana-Torrelio, William B. Karesh, Peter Daszak

## Abstract

The global wildlife trade network is a massive system that has been shown to threaten biodiversity conservation, introduce non-native species and pathogens, and cause chronic animal welfare concerns. Despite its scale and impact, comprehensive characterization of the global wildlife trade is hampered by data that are limited in their temporal or taxonomic scope and detail. To help fill this gap, we present data on 15 years of the importation of wildlife and their derived products into the United States (2000-2014), originally collected by the United States Fish and Wildlife Service. We curated and cleaned the data and added taxonomic information to improve data usability. These data include > 2 million wildlife or wildlife product shipments, representing > 60 biological classes and > 3.2 billion live organisms. These data will be broadly useful to both scientists and policymakers seeking to better understand the volume, sources, biological composition, and potential risks of the global wildlife trade.

## Background & Summary

The wildlife trade represents a major threat to the conservation of many species due to the harvest and depletion of wild populations for the purpose of trade in animals and/or their derived products ^1-6^. Consequently, understanding trade patterns and drivers is essential to mitigating the negative effects of trade on ecosystems, including those on which humanity depends ^7^. Characterization of the direct harvest and subsequent trade in wildlife is conceptually straightforward and should be aided by existing governmental monitoring programs. Currently, however, data on biological resource use are particularly scarce relative to information on other conservation threats, and the utility of existing datasets is often limited by a narrow taxonomic focus ^8^. Furthermore, comprehensive evaluation of the wildlife trade at domestic and international scales is complicated by the existence of both legal trade pathways, which are subject to differing regulations and monitoring effort in different nations, and illegal trade pathways, which are under-detected and under-reported due to their illicit nature ^9,10^. Finally, multi-country wildlife trade data sources, like the CITES Trade Database, can have reporting discrepancies and complex data structures that challenge analysis and interpretation ^11-15^. Despite these difficulties, efforts to describe and quantify the wildlife trade have scientific value, given the trade’s demonstrated impact on wildlife conservation status ^2-4,6^, animal welfare ^16^, the introduction of non-native species ^17-19^, and the spread of non-native pathogens, including zoonoses that may threaten human health ^9,10,20,21^.

The United States Fish and Wildlife Service’s (USFWS) Law Enforcement Management Information System (LEMIS) data have been used as a resource for research on the legal wildlife trade. These data, derived from legally mandated reports submitted to USFWS ^10^, contain information on US imports/exports of both live organisms and wildlife products. Previous studies, having obtained LEMIS records through Freedom of Information Act (FOIA) requests, have used the data to address broad temporal and taxonomic patterns in the US wildlife trade ^7,10^ and trends in the trade of specific focal taxa ^15,22-24^. However, the LEMIS trade data underlying analyses have either not been shared as part of the publication process, or the data that have been released focus on relatively limited time periods and study taxa. In addition, to the best of our knowledge, LEMIS data are not permanently archived ^10^, and independent parties acquiring LEMIS data may obtain subtly different datasets depending upon the date and specifics of their data requests. These factors, combined with the time investment and domain-specific knowledge required to request, process, and interpret LEMIS records, are likely barriers to the wider use of LEMIS data and may muddle comparability among studies.

Here, we collate and share 15 years of USFWS LEMIS wildlife trade importation data. While previous studies have summarized different portions of these data ^7,10,22^, the complete, cleaned dataset has not been released until now. Furthermore, we provide an R package interface for the dataset, aiming to streamline data access and ease the key initial analytical steps of data manipulation and visualization. This dataset will be of broad interest to researchers investigating the conservation impacts of overexploitation through trade, the introduction of alien species, and the potential health impact on humans, native wildlife, and domesticated species of the widespread transport of wildlife that may harbor pathogens of concern. Critically, it represents a single data resource that is relevant to researchers working across diverse taxonomic groups, allowing for greater comparability across wildlife trade work in the future.

## Methods

On a consistent basis since the mid-2000s, we have filed FOIA requests to USFWS for LEMIS data concerning importation of wildlife and wildlife products from all countries, noting that we were interested in both legal and illegal products that were documented and/or seized by US authorities. Specifically, we requested: taxonomic information (i.e., species identity or lowest-level taxonomic identification available), value of the product (reported in US dollars), wildlife description (i.e., type of wildlife product such as “live” or “skin”), quantity, unit (of the quantity metric), country of origin, country of shipment, action taken by USFWS on import, final disposition decision, date of disposition, date of shipment, the US port where the product was received, the US importer, and the foreign exporter (Table 1). At the time of writing, these requests have generated 15 years of US wildlife importation data spanning from 2000 through 2014 ^25^. As we continue to file requests for LEMIS data, the version-controlled Zenodo data repository and R package will be updated accordingly.

**Table 1.** LEMIS metadata showing data fields and field descriptions for all variables appearing in the cleaned dataset. EHA = EcoHealth Alliance, USFWS = United States Fish and Wildlife Service.

| Field | Description |
| --- | --- |
| control_number | Shipment ID number |
| species_code | USFWS code for the wildlife product |
| taxa | USFWS-derived broad taxonomic categorization |
| class | EHA-derived class-level taxonomic designation |
| genus | Genus (or higher-level taxonomic name) of the wildlife product |
| species | Species of the wildlife product |
| subspecies | Subspecies of the wildlife product |
| specific_name | A specific common name for the wildlife product |
| generic_name | A general common name for the wildlife product |
| description | Type/form of the wildlife product |
| quantity | Numeric quantity of the wildlife product |
| unit | Unit for the numeric quantity |
| value | Reported value of the wildlife product in US dollars |
| country_origin | Code for the country of origin of the wildlife product |
| country_imp_exp | Code for the country to/from which the wildlife product is shipped |
| purpose | The reason the wildlife product is being imported |
| source | The type of source within the origin country (e.g., wild, bred) |
| action | Action taken by USFWS on import ((C)leared/(R)efused) |
| disposition | Fate of the import |
| disposition_date | Full date when disposition occurred |
| disposition_year | Year when disposition occurred (derived from 'disposition_date') |
| shipment_date | Full date when the shipment arrived |
| shipment_year | Year when the shipment arrived (derived from 'shipment_date') |
| import_export | Whether the shipment is an (I)mport or (E)xport |
| port | Port or region of shipment entry |
| us_co | US party of the shipment |
| foreign_co | Foreign party of the shipment |
| cleaning_notes | Notes generated during data cleaning |

Data processing is described here only in broad outline both for brevity and because the entire data cleaning workflow is publicly available for inspection (see “Code availability” section below). Raw LEMIS data were provided by the USFWS as Microsoft Excel files, and file structure varied slightly across request responses. We aggregated these data into a single database, and performed a variety of quality assurance and data cleaning operations to improve data integrity and usability. All data processing and cleaning took place within the R statistical programming environment ^26^.

First, we harmonized data indicating missingness and other uninterpretable field values (i.e., “***”) to the standard missing data value in R (i.e., NA values). Although our data requests specified our interest in imported wildlife or wildlife products, a small proportion of the data we received (< 5%) did not contain values of “I” (indicating “import”) in the ‘import_export’ data field. Because we couldn’t confidently assess whether these records represented imported products, we removed them from the dataset. We also discovered a subset of records from one shipment year (2013) that were composed of near-duplicate records. These comprised rows that were exact duplicates of one another except for the ‘value’ field; one portion of the data for these near-duplicate matches recorded missing data for the ‘value’ field, while the other portion recorded numeric values. Given that all of the records containing missing ‘value’ data in this near-duplicate set were from the same raw data file, we deduced that we received duplicated information for this set of records, with one version of the records containing the ‘value’ data that was missing in the other. We removed the near-duplicate records that contained missing ‘value’ data, retaining the near-duplicates with good ‘value’ data.

We then cleaned data fields that should have been restricted to specific, coded values, comparing the values observed in the raw data with valid codes as indicated by USFWS code key documentation (available in our code repository). We converted irregular code entries to valid codes where it was possible to do so with reasonable confidence given the data context. In some cases, irregular code entries were apparent typographic errors. For example, in the ‘description’ field, “MEA” is the code used to indicate a meat product. We therefore assumed that records with a ‘description’ entry of “MAE” and a declared unit of kilograms were likely erroneous entries of the valid code “MEA”. In other cases, irregular codes seemed to be data entry errors resulting from subtle differences between commonly used abbreviations and the actual, valid codes for LEMIS data. For example, valid codes for the ‘unit’ field are two characters long; we thus assumed any ‘unit’ entries of “L” were meant to indicate a unit of liters, which should be expressed with the valid code “LT”. When we were unable to reasonably infer a particular data entry error, we converted irregular codes to a value of “non-standard value”. We also generated a ‘cleaning_notes’ field in the final dataset which preserves the original values that were converted to “non-standard value” for users who wish to attempt interpretation of the raw data. The following fields were cleaned in this manner: ‘description’, ‘unit’, ‘country_origin’, ‘country_imp_exp’, ‘purpose’, ‘source’, ‘action’, ‘disposition’, and ‘port’ (Table 1).

Next, we attempted to clean disposition date data. While the shipment dates in the raw data we received were strictly within the bounds of the years requested (i.e., 2000-2014), likely because this field was used by the USFWS to pull the data, the disposition date field was more varied. Some disposition date entries were obviously erroneous (e.g., those listing dates in the future) while others were likely artifacts resulting from data storage and sharing processes (e.g., when using Microsoft Excel files, blank values in date-formatted fields can sometimes be converted to unintended default date values). The vast majority of raw records in the dataset (> 95%) list a disposition date identical to or later than the shipment date. Because logically a disposition decision should occur after a product is received, where there were obvious conflicts between the shipment date and disposition date, we assumed disposition dates should refer to a date on or after the shipment date. Thus, we cleaned all obviously problematic disposition dates, particularly those lying outside the time period 2000-2014. Note, however, that disposition dates in 2015 may be sensible and valid for shipments received late in 2014.

Next, we cleaned and supplemented taxonomic information in the LEMIS data. Using the provided ‘species_code’ field and USFWS keys, we were able to derive a ‘taxa’ field for the vast majority (> 97%) of records (Table 1). However, this USFWS-defined ‘taxa’ categorization, while useful for general data inspection, does not correspond to a consistent taxonomic concept. Therefore, we sought to designate a taxonomic class for all LEMIS data where possible. We used the R package ‘taxadb’ to automatically gather class information ^27^, drawing primarily from the taxonomic classification provided by the Catalogue of Life (COL) database. Where the COL data did not allow for automated class-level taxonomic calls, we drew from the Integrated Taxonomic Information System (ITIS), harmonizing data with the COL class categorization. Furthermore, the lack of automatic class-level taxonomic assignment for some taxonomic entries alerted us to raw values potentially in need of correction, initiating an iterative data cleaning process. First, as part of this cleaning, vague or missing taxonomic information in the ‘species’ and ‘subspecies’ fields were converted to “sp.” values for consistency. Next, we manually inspected and corrected unique combinations of the ‘genus’, ‘species’, ‘subspecies’, ‘specific_name’, and ‘generic_name’ fields (Table 1). In many cases, errors represented minor misspellings (e.g., *Philetarius socius* instead of *Philetairus socius*) or inversions of the genus and species names. Finally, where we were still unable to recover automated class-level information, we manually assigned class when data specificity and context from other fields allowed. Many of these data represented cases where the LEMIS data uses alternate taxonomy that is not recognized by either the COL or the ITIS. Nonetheless, the data provided often enabled unambiguous class-level assignment.

### Code availability

Our custom R package, which provides access to the data described here, is publicly available at https://github.com/ecohealthalliance/lemis. Installation of the package and subsequent download of the data enables efficient, on-disk manipulation of the entire cleaned dataset ^28,29^. Basic package usage is outlined in the main package README file on the GitHub site. The code implementation of the data cleaning process described above is also available in the package codebase (via the ‘data-raw’ directory) and is outlined in the associated developer README file. These scripts span the entirety of our data processing and cleaning workflow, from importation and collation of the raw USFWS LEMIS data files through to generation of the single, cleaned data file as discussed in this manuscript. Thus, the scripts serve as transparent, reproducible documentation of our data processing in full.

## Data Records

We present over > 5.5 million USFWS LEMIS wildlife or wildlife product records spanning 15 years and 28 data fields ^25^. These records were derived from > 2 million unique shipments processed by USFWS during the time period and represent > 3.2 billion live organisms (Fig. 1). We provide the final cleaned data as a single comma-separated value file. Original raw data as provided by the USFWS are also available in the data repository. Although relatively large (∼1 gigabyte), the cleaned data file can be imported into a software environment of choice for data analysis. Alternatively, our R package provides access to a release of the same cleaned dataset but with a data download and manipulation framework that is designed to work well with this large dataset, as previously described.

**Figure 1.**
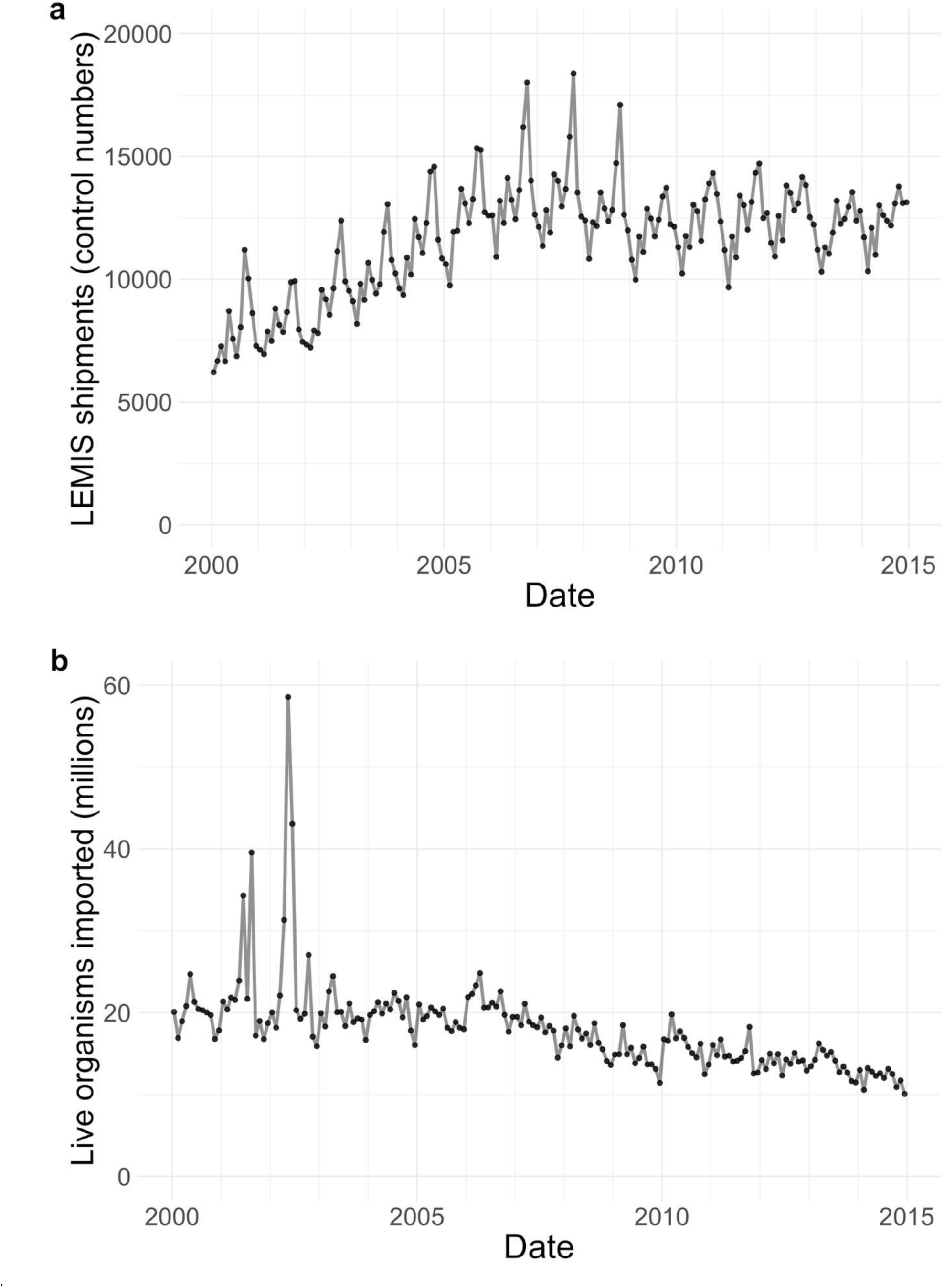
Number of unique shipments (a) and number of live organisms (b) imported per month in the LEMIS wildlife trade data from 2000 through 2014. We defined shipments as synonymous with the LEMIS data field ‘control_number’. Each shipment may contain multiple types of wildlife products and thus be recorded over multiple rows in the data. Note that the spikes in live organism imports in 2001 and 2002 are driven by extremely large recorded shipments (> 5 million individuals) of tropical fish and crustaceans (*Penaeus* sp.).

Twenty-three of the final data fields are cleaned versions of the original data provided by the USFWS: ‘control_number’, ‘species_code’, ‘genus’, ‘species’, ‘subspecies’, ‘specific_name’, ‘generic_name’, ‘description’, ‘quantity’, ‘unit’, ‘value’, ‘country_origin’, ‘country_imp_exp’, ‘purpose’, ‘source’, ‘action’, ‘disposition’, ‘disposition_date’, ‘shipment_date’, ‘import_export’, ‘port’, ‘us_co’, and ‘foreign_co’ (Table 1). To these original data fields, we added five: ‘taxa’, ‘class’, and ‘cleaning_notes’ (all as previously described), as well as ‘dispostion_year’ and ‘shipment_year’ (derived from ‘disposition_date’ and ‘shipment_date’, respectively). To briefly describe the LEMIS data fields, we consider ‘control_number’ to represent a unique individual shipment processed by the USFWS (Fig. 1). Different wildlife products contained within the same shipment may be represented in the LEMIS data by multiple data rows, all of which share a common ‘control_number’. Consistent with this interpretation, all rows of data sharing the same ‘control_number’ share the same country of shipment and shipment date. Different products within the same shipment may differ in other ways, however. For example, they may have been originally derived from different countries and may have different disposition histories. Next, the ‘species_code’, ‘taxa’, ‘class’, ‘genus’, ‘species’, ‘subspecies’, ‘specific_name’, and ‘generic_name’ columns all provide information serving to identify the wildlife or wildlife product (Table 1). While the ‘genus’ column largely corresponds to taxonomic genus, sometimes higher-level categorizations were provided in this field, apparently when the genus was unknown. Using our automated taxonomic calling workflow, we were able to assign ‘class’ information to > 92% of LEMIS records. All further data fields besides ‘cleaning_notes’ serve to detail the wildlife product, as outlined in Table 1. Note that although we consistently requested product ‘value’ information from the USFWS, it was not provided for four years of LEMIS data (2008-2010 and 2014).

## Technical Validation

Following data cleaning, which primarily aimed to ensure that all relevant data fields contained valid USFWS-defined codes, we validated our final dataset by plotting the distribution of unique values and value string lengths across all data fields. These checks serve to verify that fields only contain expected values/codes and that the string length of entries in free text fields (e.g., ‘genus’, ‘species’) were not abnormally short or long, which could indicate problematic entries.

## Usage Notes

While we did remove what we believe to be erroneous near-duplicate records in the dataset (as described in the Methods), end users should note that exact duplicate records remain. This is because even exact duplicate records may represent accurate data, especially in cases where the recorded ‘quantity’ value is 1. For example, in the final dataset, ‘control_number’ 2000732392 records the importation of a shipment of garments from France which were themselves derived from reticulated pythons (*Python reticulatus*) originating in Malaysia. Within this ‘control_number’ value (representing one shipment), one data record, reporting a ‘quantity’ of 1 and a ‘value’ of $1,458, is duplicated 25 times. Our assumption is that these garments, and similar duplicate products, were individually packaged but shipped together such that officers at the port of entry recorded exact duplicate data entries to capture the total product volume within the shipment. In other cases, similar information may have been aggregated during data entry (e.g., recording the identical product data as a single record with a quantity of 25). We verified that all duplicate records that remain in the data originated from the same raw data file. This indicates that these records were provided as such by USFWS and ensures they were not artifacts generated through our data processing pipeline (e.g., by combining data across multiple raw data files that contained overlapping information). Thus, we believe we have made the most conservative data processing decision by preserving the original form of the data unless we had good reason to perform data cleaning. Nevertheless, users should be aware of the potential presence of duplicate records in any data subset of interest, and these records should be scrutinized for inclusion in analyses given the specific study objectives.

The dataset provides multiple, complementary data fields reporting taxonomic identity that deserve special attention. Generally, users will want to consider the ‘taxa’ and ‘class’ fields in conjunction to analyze trade data for large taxonomic groups. While ‘class’ is typically a more specific taxonomic designation, ‘taxa’ has fewer missing values in the final dataset (‘class’ information available for > 92% of LEMIS records; ‘taxa’ information available for > 97% of LEMIS records). Which field deserves greater focus will depend on the analytical goals. For example, the ‘taxa’ category “fish” encompasses LEMIS records representing six distinct ‘class’ values: Actinopterygii, Cephalaspidomorphi, Elasmobranchii, Holocephali, Myxini, and Sarcopterygii. Clearly, ‘class’ is biologically meaningful and may help users rapidly narrow their analytical focus, but users should keep in mind that there are records within the ‘taxa’ category of “fish” for which ‘class’ could not be unambiguously assigned. For some research questions, these data may also be of interest.

In addition, users must be cognizant of the fact that taxa may be represented by multiple taxonomic synonyms. While we sought to provide high-level taxonomic information (e.g., class assignments) that would help users in generating a relevant data subset for analysis, we did not attempt to synonymize species-level names given the large number of taxa present in the LEMIS data and the constantly shifting (and contentious) landscape of preferred taxonomic nomenclature. End users will need to apply their expertise on taxa of interest in order to generate sound taxonomic delineations where synonymies exist in the data.

Furthermore, data users should be cautious about their interpretation of the ‘shipment_date’ and ‘disposition_date’ fields. As previously mentioned, while ‘shipment_date’ entries within the raw data we received fell completely within the time period of 2000-2014, ‘disposition_date’ ranged more widely. Even following data cleaning to harmonize ‘disposition_date’ entries that were obviously problematic, significant discrepancies between ‘shipment_date’ and ‘disposition_date’ still exist for some records in the final dataset. We have chosen to preserve these data as is there is no clear cut-off at which differences between disposition date and shipment date become invalid. For example, dispositions that occur months after the declared shipment date could reflect the reality of product processing even though a large majority of records (> 70%) indicate that disposition typically occurs within a week of the shipment date. Certainly, users should be wary of any disposition date values that precede the associated shipment date, as we are unaware how this could represent an accurate accounting of the product disposition process. However, for many potential analyses, differences in the date fields may not be a significant cause for concern because ‘shipment_date’ alone provides a sound index for those interested in temporal trends in wildlife trade.

Finally, data users should be careful about interpreting the ‘country_imp_exp’ and ‘country_origin’ data fields. These fields are meant to represent the most recent location (‘country_imp_exp’) and point of origin (‘country_origin’) for the wildlife or wildlife products, but data in these fields are derived from import documents completed by the importer and are therefore not verifiable. Complex import/export histories can result in surprising entries for these fields ^21^. For example, rodents of the genus *Abrocoma* are native to South America. However, our data describe a shipment of garments derived from *Abrocoma* sp. (‘control_number’ 2008273877) with a ‘country_imp_exp’ of Switzerland and a ‘country_origin’ of Hungary. The apparent contradiction in this case is resolved by recognizing that the ‘source’ column indicates these animals were derived from a domestic ranching operation rather than being taken directly from the wild. However, for those interested in the true origins of wildlife and wildlife products that are sourced from the wild (∼78% of our data records), the ‘country_origin’ field deserves special scrutiny to ensure the recorded country is in fact a biologically-realistic point of origin for the species in question.

Understanding the appropriate interpretation of the ‘country_imp_exp’ and ‘country_origin’ fields also illuminates how seemingly incongruous records listing the US as the ‘country_origin’ for a US importation can in fact be valid data. For example, ‘control_number’ 2005537093 represents a shipment of shoe products derived from white-tailed deer (*Odocoileus virginianus*). The ‘country_origin’ is recorded as the US, where the wildlife was presumably originally harvested, while Italy is recorded as the ‘country_imp_exp’ since this was the proximate source of the shoe products. Hence, for wildlife products where some part of the manufacturing process takes place abroad, it is indeed expected that raw materials derived from US wildlife are shipped internationally, thereby resulting in LEMIS data that indicate the US importation of a wildlife product that was originally sourced from the US.

## Acknowledgements

The authors wish to express their sincere thanks to the United States Fish and Wildlife Service and the numerous employees whose prompt, professional service over the years has helped make this data more widely available to the scientific community. The work in this paper was supported by: a National Science Foundation Human and Social Dynamics ‘Agents of Change’ award (SES-HSD-AOC “Human-Related Factors Affecting Emerging Infectious Diseases”, BCS-0826779 and BCS-0826840); a National Institutes of Health NIGMS grant (1R01GM100471-01, “MASpread”); a Joint NSF-NIH-USDA/BBSRC Ecology and Evolution of Infectious Diseases award (NSF DEB 1414374, BBSRC BB/M008894/1, “US-UK Collab: Risks of Animal and Plant Infectious Diseases through Trade (RAPID Trade)”); the United States Agency for International Development (USAID) Emerging Pandemic Threats PREDICT project; and core funding from EcoHealth Alliance.

## Author Contributions

K.M.S., A.M.W., K.F.S., J.P.R., C.Z.-T., W.B.K., and P.D. designed, drafted, and filed Freedom of Information Act requests. E.A.E., A.M.W., and C.Z-T. made key contributions to the LEMIS data processing and cleaning workflow. N.R. developed and maintains the R package for data access. E.A.E. drafted the manuscript, and all authors were involved in editing and approving the final manuscript.

